# Reciprocal regulation of TLR4, TLR3 and Macrophage Scavenger Receptor 1 regulates nonopsonic phagocytosis of the fungal pathogen *Cryptococcus neoformans*

**DOI:** 10.1101/2023.01.30.525903

**Authors:** Chinaemerem U. Onyishi, Guillaume E. Desanti, Alex L. Wilkinson, Gyorgy Fejer, Olivier D. Christophe, Clare E. Bryant, Subhankar Mukhopadhyay, Siamon Gordon, Robin C. May

**Affiliations:** Institute of Microbiology & Infection and School of Biosciences, University of Birmingham, Edgbaston, Birmingham, B15 2TT, United Kingdom; School of Biomedical Sciences, Faculty of Health, University of Plymouth, Plymouth, United Kingdom; Laboratory for Hemostasis, Inflammation & Thrombosis, Unité Mixed de Recherche 1176, Institut National de la Santé et de la Recherche Médicale, Université Paris-Saclay, 94276 Le Kremlin-Bicêtre, France; University of Cambridge, Department of Medicine, Box 157, Level 5, Addenbrooke’s Hospital, Hills Road, Cambridge, CB2 0QQ, United Kingdom; Peter Gorer Department of Immunobiology, School of Immunology & Microbial Sciences, King’s College London, SE1 9RT, United Kingdom; Department of Microbiology and Immunology, College of Medicine, Chang Gung University, Taoyuan, Taiwan; Sir William Dunn School of Pathology, University of Oxford, Oxford, UK

## Abstract

The opportunistic fungal pathogen *Cryptococcus neoformans* causes lethal infections in immunocompromised patients. Macrophages are central to the host response to cryptococci; however, it is unclear how *C. neoformans* is recognized and phagocytosed by macrophages. Here we investigate the role of TLR4 in the nonopsonic phagocytosis of *C. neoformans*. We find that loss of TLR4 function unexpectedly increases phagocytosis of nonopsonized cryptococci. The increased phagocytosis observed in *Tlr4*^*-/-*^ cells was dampened by pre-treatment of macrophages with either a TLR3 inhibitor or oxidised-LDL, a known ligand of scavenger receptors. The scavenger receptor, macrophage scavenger receptor 1 (MSR1) (also known as SR-A1 or CD204) was upregulated in *Tlr4*^*-/-*^ macrophages and there was a 75% decrease in phagocytosis of nonopsonized cryptococci by *Msr1*^*-/-*^ macrophages. Furthermore, immunofluorescence imaging revealed colocalization of MSR1 and internalised cryptococci. Together, these results identify MSR1 as a key receptor for the phagocytosis of nonopsonized *C. neoformans* and demonstrate TLR4/MSR1 crosstalk in the phagocytosis of *C. neoformans*.

## Introduction

*Cryptococcus neoformans* is an encapsulated yeast that causes life-threatening infections in humans and other animals^1,2^, with an estimated global burden of 181,000 deaths annually^3^. Infection with *C. neoformans* begins with the inhalation of fungal cells from the environment into the lungs^1^. Within the lungs, tissue-resident macrophages are amongst the first immune cells the fungi encounter^4^, thus, the interaction between host macrophages and invading fungi is believed to be an important determinant of disease progression and outcome. Nonopsonized cryptococci are phagocytosed poorly^5^, but since opsonising antibodies are negligible within the healthy lung^6^, this low level of nonopsonic uptake is likely a critical determinant of the subsequent course of an infection. However, there is no clear understanding of the mechanism by which macrophages detect and phagocytose *C. neoformans* in the absence of opsonins^4,7^.

Phagocytosis, defined as the uptake of particles greater than 0.5 µm, is a significant process in the innate immune response as it leads to the degradation of invading pathogens and the presentation of microbial ligands on MHC molecules, thereby activating the adaptive arm of the immune system^8^. Nonopsonic phagocytosis is initiated by the recognition of pathogen associated molecular patterns (PAMPs) on the surface of microbes by host pattern recognition receptors (PRRs)^8^. PRRs on professional phagocytes include members of the Toll-like receptor (TLR) family, the C-type lectin receptor (CLR) family, and the scavenger receptor (SR) family. All of these have been implicated in the recognition of *C. neoformans* to varying degrees, with β-1,3-glucans, mannans and glucuronoxylomannan (GXM) found on the *C. neoformans* cell wall and capsule serving as PAMPs^9–15^. The CLR, Dectin-1 (also known as CLEC7A), is well-known for its role in the recognition of fungal β-glucans^11^ and has been identified as the key PRR involved in the phagocytosis of *Candida albicans*^16,17^. However, previous work found that Dectin-1 is only marginally involved in the phagocytosis of nonopsonized *C. neoformans*^5^, suggesting that other nonopsonic receptors for *C. neoformans* may be more important.

Within the TLR family, TLR4 is known to recognise fungal mannans^12^ and GXM^9^, leading to the activation of downstream signalling cascades. TLR4 signalling is mediated by the adaptor proteins myeloid differentiation primary response 88 (MyD88) and TIR-domain-containing adapter-inducing interferon-β (TRIF)^18^. The MyD88-dependent pathway is used by all TLRs except TLR3, which uses TRIF-dependent signalling instead^19,20^. The MyD88-dependent pathway and the TRIF-dependent pathway ultimately lead to the activation of the transcription factor nuclear factor κB (NF-κB) and mitogen activated protein kinases (MAPKs)^20^. The TRIF pathway also leads to the activation of Interferon regulatory factor 3 (IRF3). These then act to activate the expression and secretion of proinflammatory cytokines (MyD88 and TRIF pathway) and Type I interferons (TRIF pathway)^8,18,21^. Notably, plasma membrane TLRs also activate Rap GTPase and Rac GTPase to activate phagocytic integrins and other bona fide phagocytic receptors which are then responsible for pathogen engulfment^22^. One prominent example is seen in the collaboration between TLR2 and Dectin-1 in modulating the uptake of β-glucan and subsequent cytokine production^23,24^. Similarly, there are numerous examples of TLR crosstalk with SRs^25,26^.

Whilst investigating the role of TLR signalling in the inflammatory response to cryptococci, we made the unexpected discovery that loss of TLR4 activity leads to enhanced nonopsonic uptake of the fungus. We show that this increase was driven by crosstalk between TLR4 and TLR3 to regulate the surface expression of Macrophage Scavenger Receptor 1 (MSR1) (also known as SR-A1 or CD204), such that the loss of TLR4 signalling increased the expression of MSR1, but not other SRs, leading to increased uptake. We provide evidence that MSR1 is a necessary receptor for the nonopsonic phagocytosis of *C. neoformans*, shedding light on a key host receptor involved in the uptake of this fungal pathogen.

## Results

### Both chemical inhibition and genetic loss of TLR4 signalling results in an increase in the phagocytosis of nonopsonized *C. neoformans*

To investigate the role of TLR4 on the phagocytosis of nonopsonized *C. neoformans*, J774A.1 murine macrophages were treated with 0.2 μM TAK-242, an inhibitor of TLR4 signalling, for 1 h before being infected with *C. neoformans* still in the presence of the inhibitor. Surprisingly, TLR4 inhibition resulted in a significant 1.7-fold increase in the phagocytosis of nonopsonized cryptococci (Figure 1A). We tested whether genetic loss of *TLR4* would replicate this effect by using immortalised bone marrow derived macrophages (iBMDMs) isolated from wildtype and *Tlr4*^*-/-*^ C57BL/6 mice. As with the chemical inhibition of TLR4, genetic knockout of *TLR4* led to a pronounced 8-fold increase in the phagocytosis of nonopsonized *C. neoformans* (Figure 1B and C).

**Figure 1:**
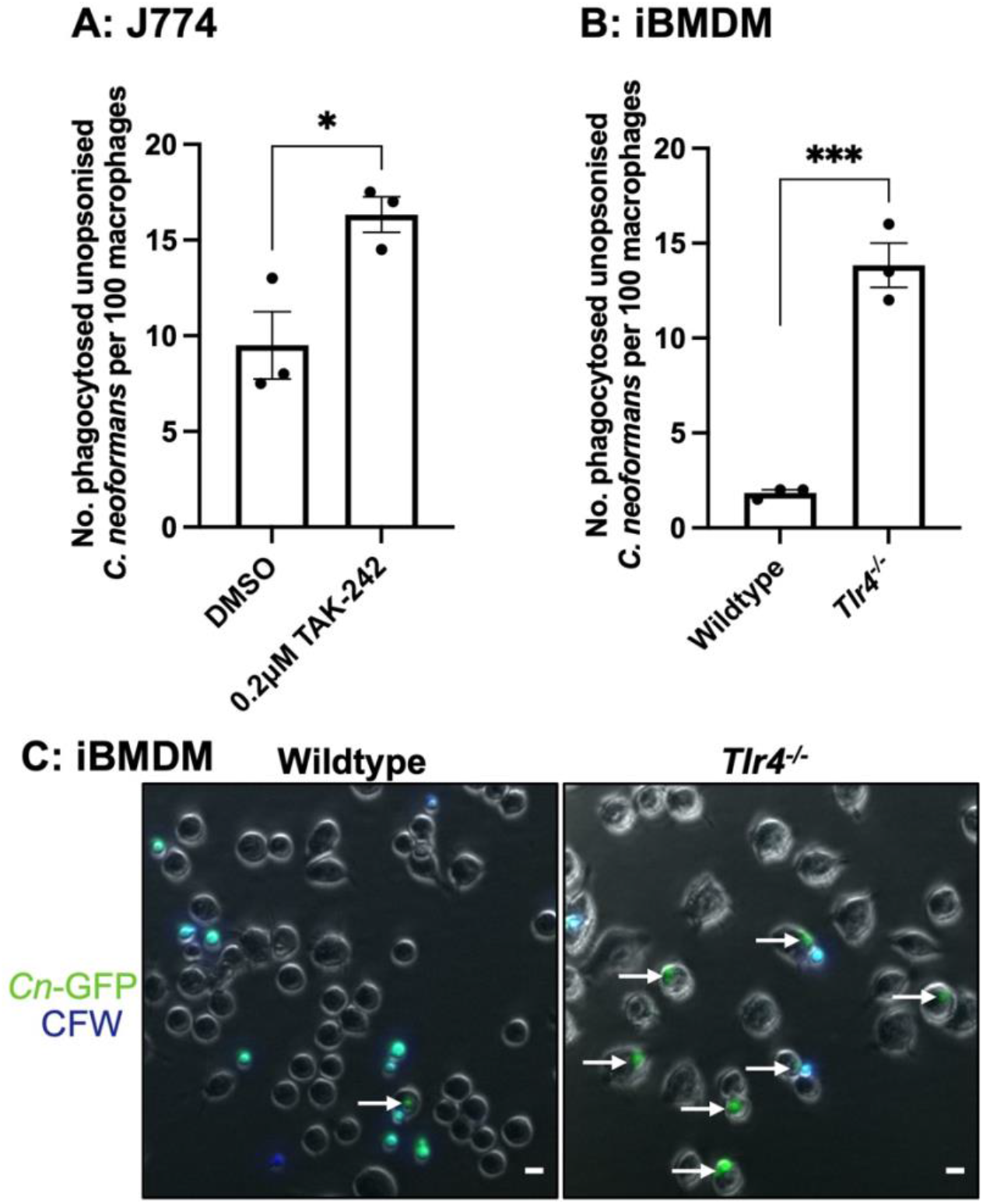
Both chemical inhibition and genetic loss of TLR4 results in an increase in the phagocytosis of *C. neoformans*. **(A)** J774A.1 macrophages were treated with DMSO (control) or 0.2 μM TAK-242, a TLR4 specific inhibitor, for 1 h before infection with nonopsonized *C. neoformans*. **(B)** Immortalised bone marrow derived macrophages (iBMDM) from wildtype and *Tlr4*^*-/-*^ macrophages were infected with nonopsonized *C. neoformans*. Phagocytosis was quantified as the number of individual internalised cryptococci within 100 macrophages. Figures are representative of at least three independent technical repeats. Data shown as mean ± SEM; each data point represents one biological replicate for each condition; statistical significance was evaluated using a t-test: *p<0.05, ***p<0.001. **(C)** Representative image showing the phagocytosis of GFP-labelled *C. neoformans* (*Cn*-GFP) by wildtype and *Tlr4*^*-/-*^ iBMDM. Calcofluor White (CFW) was used to stain extracellular fungi. White arrows show phagocytosed fungi. Scale bar = 10 μm.

### Increased uptake observed in *Tlr4*^*-/-*^ macrophages is not a consequence of increased intracellular proliferation

Proinflammatory responses, such as those driven by TLR4, have been shown to restrict the intracellular proliferation of cryptococci^27^, hence we considered that the perceived increase in phagocytosis might instead reflect increased proliferation in the absence of TLR4 activity. To test this hypothesis, we conducted live imaging of infected macrophages and quantified the number of internalised fungi at the beginning of the video (T0) and 10 h post infection (T10) to determine the intracellular proliferation rate (IPR). This time-lapse-based IPR assay revealed that neither TLR4 inhibition using TAK-242 (Figure 2A) or *TLR4* knockout (Figure 2B) altered the IPR of cryptococci compared to control macrophages. This suggests that the observed intracellular burden of *C. neoformans* is representative of the initial rate of uptake and not due to differences in the subsequent proliferation of the fungi within macrophages.

**Figure 2:**
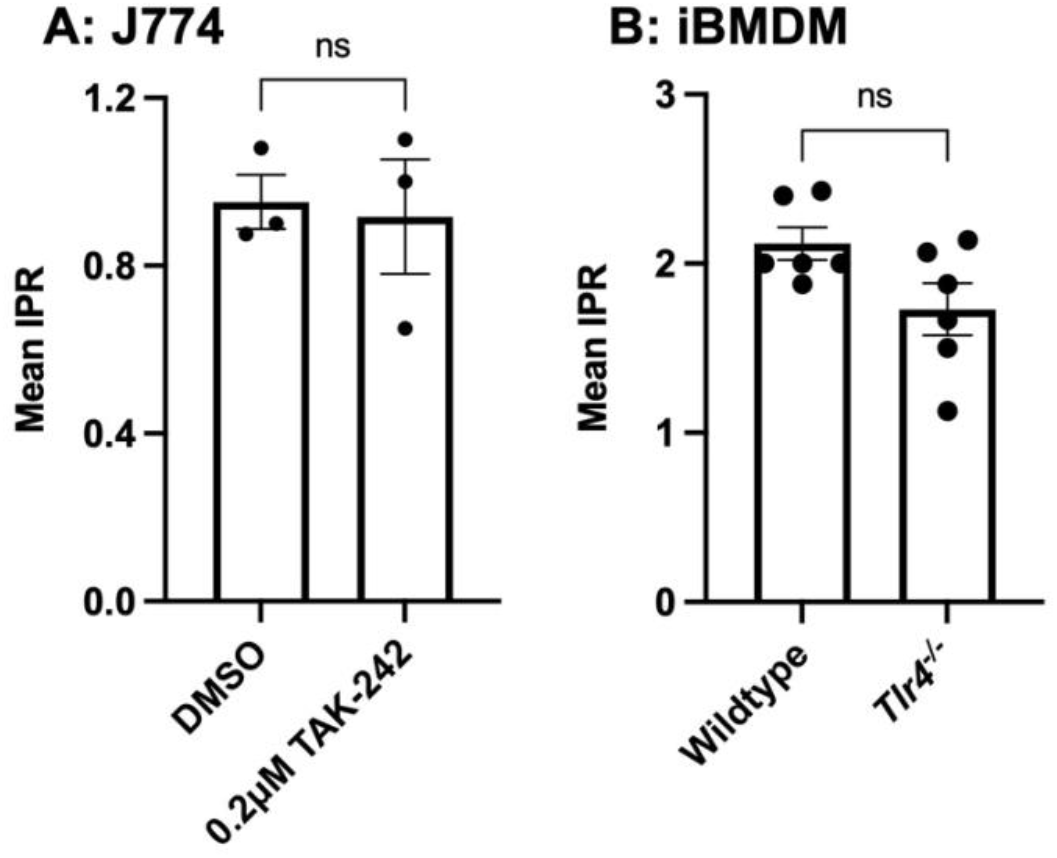
Intracellular Proliferation Rate (IPR) of *C. neoformans* within macrophages. The intracellular proliferation of *C. neoformans* was measured in **(A)** J774A.1 macrophages treated with DMSO (control) or 0.2 μM TAK-242, a TLR4 specific inhibitor, for 1 h prior to infection and **(B)** wildtype and *Tlr4*^*-/-*^ iBMDMs. After phagocytosis, extracellular *C. neoformans* was washed off and macrophages were imaged every 5 mins for 18 h. The number of internalised fungi per 100 macrophages at the ‘first frame’ (T0) and ‘last frame’ (T10) was quantified and IPR was determined using the equation: IPR = T10/T0. Data shown as mean ± SEM; ns, not significant in a t-test. The iBMDM data is pooled from two independent technical repeats performed in triplicates.

### The increased uptake observed in *Tlr4*^*-/-*^ macrophages is partially driven by TLR3 signalling and is dependent on MyD88 and TRIF

There is evidence of TLR-TLR crosstalk modulating cytokine expression^28^. Consequently, we wondered whether TLR-TLR crosstalk may also influence phagocytosis such that a loss of TLR4 signalling may lead to increased signalling through other, phagocytosis-promoting, TLRs. To explore the existence of such TLR-TLR crosstalk, we treated wildtype and *Tlr4*^*-/-*^ iBMDMs with inhibitors of TLR2 (CU CPT22), TLR3 (TLR3/dsRNA complex inhibitor), and TLR9 (ODN 2088) prior to infection with *C. neoformans*. These inhibitors were chosen because only TLR4, TLR2, TLR9, and TLR3 have been studied previously in the context of *C. neoformans* infection^29–31^.

Although inhibition of TLR2 and TLR9 had no impact on the enhanced phagocytosis seen in *Tlr4*^*-/-*^ macrophages, there was a 42% decrease in the number of phagocytosed fungi following TLR3 inhibition (Figure 3A). Notably, TLR3 inhibition consistently decreased the phagocytosis of cryptococcus by wildtype macrophages too, but the very low baseline uptake meant that this result was not statistically significant (Figure 3A). Interestingly, when macrophages were infected with *C. neoformans* opsonised with the anti-capsular 18B7 antibody, *TLR4*-deficiency resulted in an increase in uptake, but the effect of TLR3 inhibition was lost in both wildtype and *Tlr4*^*-/-*^ cells (Figure 3B). Therefore, the role of TLR3 in modulating the phagocytosis of *C. neoformans* is specific to nonopsonic uptake.

**Figure 3:**
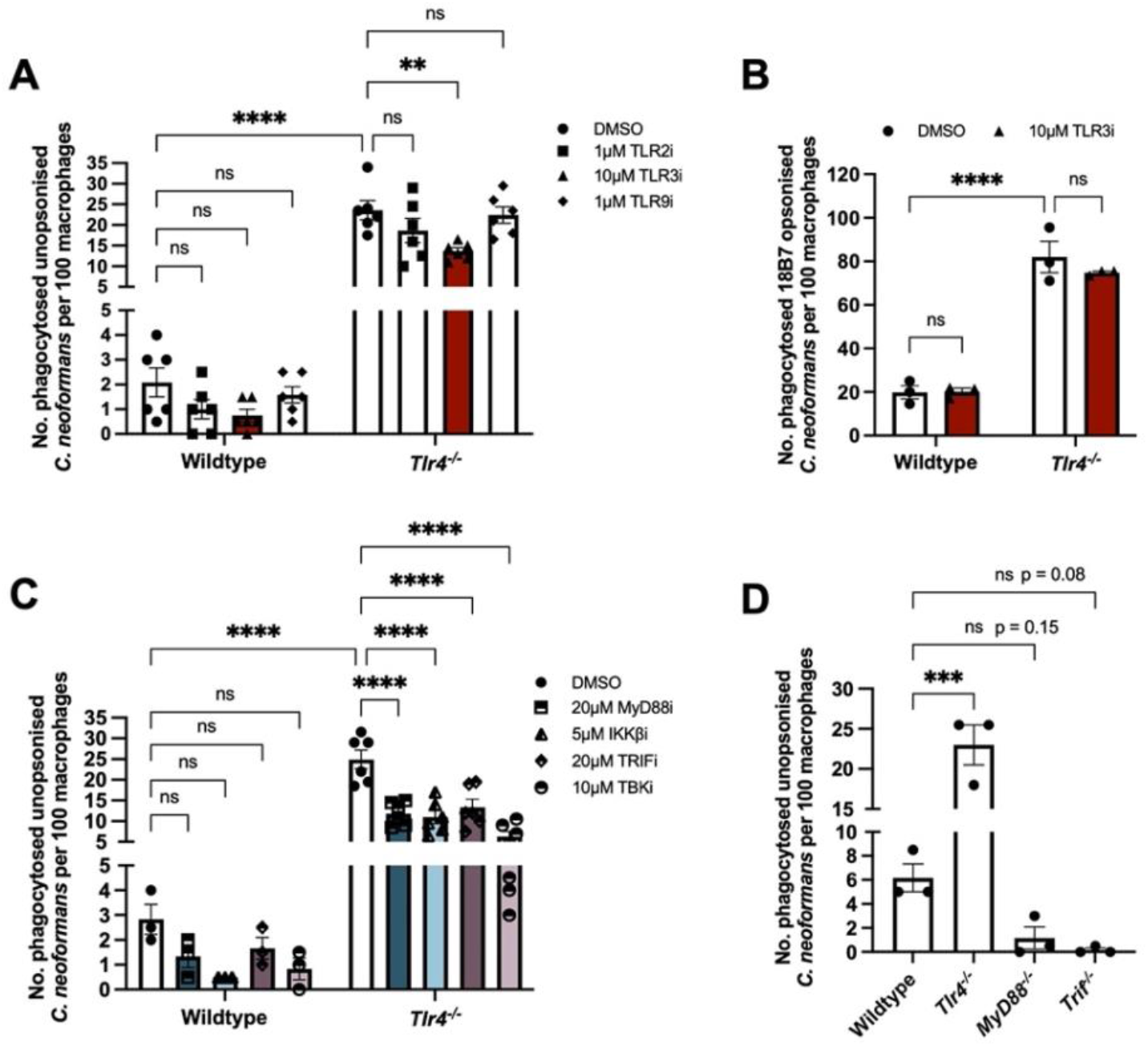
The increased phagocytosis observed in *Tlr4*^*-/-*^ macrophages is dependent on TLR3, MyD88 and TRIF signalling. **(A)** Wildtype and *Tlr4*^*-/-*^ iBMDMs were treated with chemical inhibitors of TLR2, TLR3 and TLR9 for 1 h, then infected with nonopsonized *C. neoformans*. Data is pooled from two technical repeats performed in triplicates. **(B)** Wildtype and *Tlr4*^*-/-*^ iBMDMs were treated with a TLR3 inhibitor then infected with *C. neoformans* opsonized with the anti-capsular 18B7 antibody. **(C)** Wildtype and *Tlr4*^*-/-*^ macrophages were treated with inhibitors of MyD88, IKKβ, TRIF, and TBK1. Phagocytosis was quantified as the number of internalised cryptococcus within 100 macrophages. Data shown as mean ± SEM; ns, not significant, *p<0.05, **p<0.01, ***p<0.001, ****p<0.0001 in a two-way analysis of variance (ANOVA) followed by Tukey’s post-hoc test. **(D)** Immortalised BMDMs from wildtype, *Tlr4*^*-/-*^, *MyD88*^*-/-*^ and *Trif*^*-/-*^ macrophages were infected with nonopsonized *C. neoformans*. Data is representative of at least three technical repeats. Data shown as mean ± SEM and analysed using one-way ANOVA followed by Tukey’s post-hoc test. ns, not significant, *p<0.05, **p<0.01, ***p<0.001.

TLR4 and TLR3 signalling requires the downstream adaptor molecules MyD88 (used by TLR4) and TRIF (used by both TLR4 and TLR3)^20^. Therefore, to understand the downstream signalling pathway(s) involved, macrophages were exposed to inhibitors of MyD88, TRIF, IKKβ (a kinase downstream of MyD88 that is necessary for NF-κB activation^32^), and TBK1 (a kinase downstream of TRIF that phosphorylates and activates IRF3^33^). We found that treatment with all four inhibitors dampened the increased phagocytosis observed in *TLR4*-deficient macrophages (Figure 3C). Wildtype macrophages also showed a trend towards decreased phagocytosis when treated with all four inhibitors. In line with the findings from the inhibitor treatments, macrophages derived from *MyD88*^*-/-*^ and *Trif*^*-/-*^ mice were significantly impaired in the phagocytosis of *C. neoformans* (Figure 3D).

To ensure that the significant loss of uptake in *MyD88*- and *TRIF*-deficient macrophages was not caused by an inherent deficiency in phagocytic capacity, we infected iBMDMs with CAF2-dTomato *Candida albicans*^34^. We found that *MyD88*^*-/-*^ and *Trif*^*-/-*^ macrophages had the same level of phagocytosis as wildtype macrophages (Supplementary Figure 1). Thus, non-TLR dependent pathways such as the Dectin-1 receptor that is recognised as the key PRR involved in the phagocytosis of *C. albicans*^16,17^ remain intact in *MyD88*^*-/-*^ and *Trif*^*-/-*^ macrophages. Notably, however, the loss of TLR4 also led to an increase in the phagocytosis of *C. albicans* (Supplementary Figure 1), suggesting the existence of some shared host response to both fungi. Overall, the phagocytosis of nonopsonized *C. neoformans*, but not *C. albicans*, is dependent on MyD88 and TRIF.

### Oxidised low-density lipoprotein (ox-LDL) competitively inhibits the phagocytosis of nonopsonized *C. neoformans*

Although we have shown that *TLR4*-deficiency increases the phagocytosis of nonopsonized *C. neoformans* through crosstalk with TLR3 in a MyD88- and TRIF-dependent manner, the plasma membrane receptor responsible for this increase in uptake remains unknown. Plasma membrane TLRs are not bona fide phagocytic receptors, since they are not directly responsible for the engulfment of whole microorganisms^22^. Consequently, we considered whether instead they may be modulating the availability of one or more phagocytic receptors that bind nonopsonized *C. neoformans*.

Scavenger receptors, a family of receptors that were initially identified for their role in the uptake of modified host lipoproteins^35^, are increasingly being implicated as receptors for a variety of microbes and their ligands^36–38^. Moreover, it has been shown that the expression of several scavenger receptors is upregulated in *Tlr4*^*-/-*^ mice^39^ and that TLR agonists increase the phagocytosis of *Escherichia coli* by inducing the expression of scavenger receptors^40^. We therefore tested whether the loss of TLR4 signalling increases the phagocytosis of nonopsonized *C. neoformans* through the upregulation of scavenger receptors.

Firstly, we treated macrophages with ox-LDL, a general scavenger receptor ligand and competitive inhibitor, prior to infection with *C. neoformans*. We found that ox-LDL was able to competitively inhibit the phagocytosis of *C. neoformans* in both wildtype and *Tlr4*^*-/-*^ macrophages (Figure 4A and B). When macrophages were infected with 18B7 antibody-opsonized fungi to drive uptake through Fcγ-receptors instead, ox-LDL pre-treatment had no impact on the phagocytosis of cryptococci in both wildtype and *Tlr4*^*-/-*^ macrophages (Figure 4C), indicating that the inhibition is specific to nonopsonic uptake.

**Figure 4:**
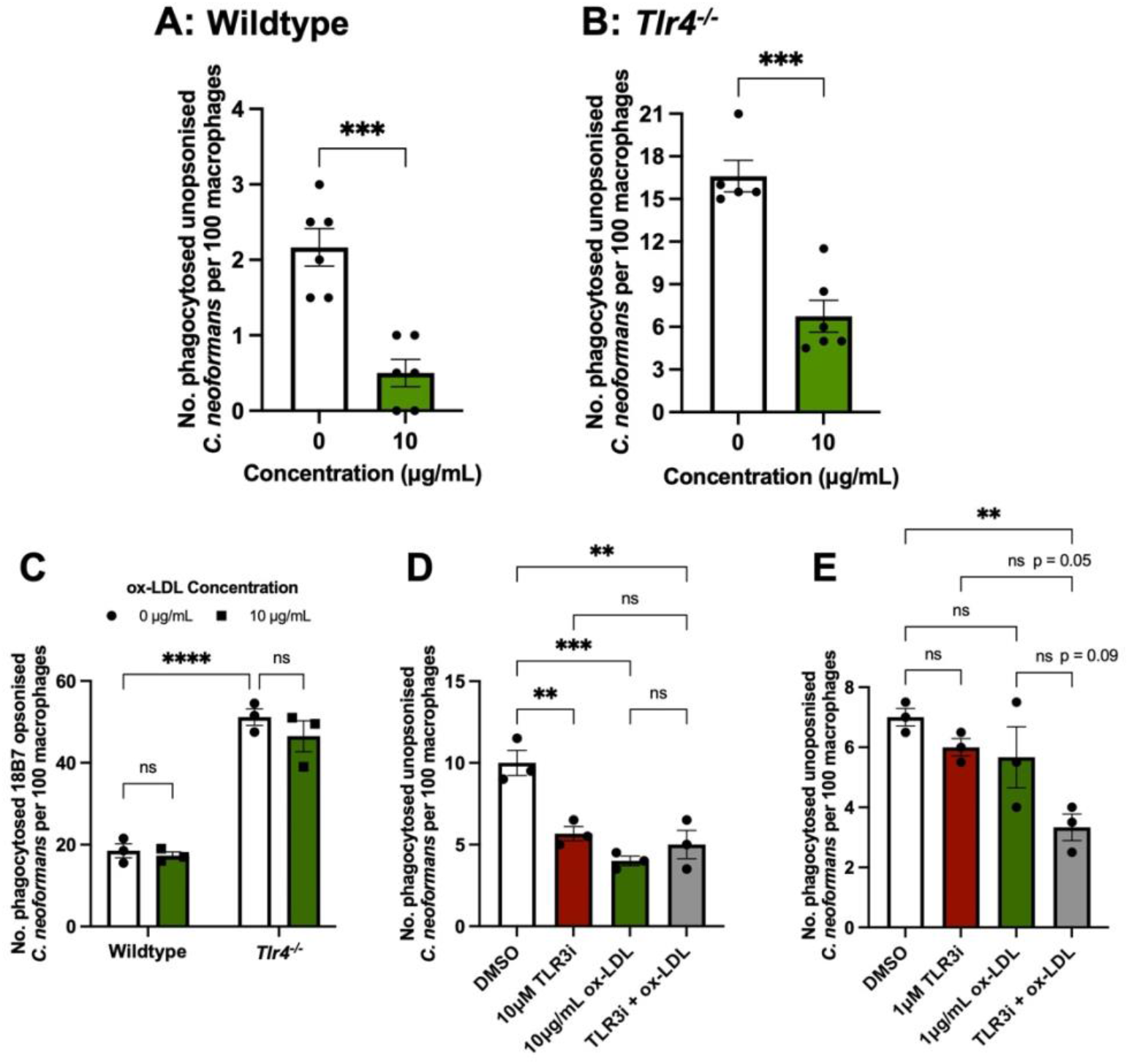
The increased phagocytosis observed in *Tlr4*^*-/-*^ macrophages is partially driven by scavenger receptors. **(A)** Wildtype and **(B)** *Tlr4*^*-/-*^ iBMDMs were treated oxidised low-density lipoprotein (ox-LDL), a general scavenger receptor ligand, for 30 mins prior to infection with nonopsonized *C. neoformans*. Data is pooled from two technical repeats. **(C)** Wildtype and *Tlr4*^*-/-*^ iBMDMs were treated with 10 μg/mL ox-LDL for 30 mins, then infected with *C. neoformans* opsonized with the anti-capsular 18B7 antibody. **(D, E)** *Tlr4*^*-/-*^ macrophages were pre-treated with optimal and suboptimal concentrations of TLR3 inhibitor and ox-LDL individually and in combination. The number of internalised fungi per 100 macrophages was quantified from fluorescent microscopy images. Data is representative of two technical repeats. Data shown as mean ± SEM; ns, not significant, *p<0.05, **p<0.01, ***p<0.001, ****p<0.0001 in a t-test **(A, B)**; two-way ANOVA **(C)**; or one-way ANOVA **(D, E)** followed by Tukey’s post-hoc test.

### TLR3 and Scavenger Receptors act in synergy along the same pathway to modulate phagocytosis

The data presented above suggest a model in which loss of TLR4 triggers increased TLR3 signalling, leading to upregulation of scavenger receptors. To test this, *Tlr4*^*-/-*^ iBMDMs were pre-treated with TLR3i and ox-LDL individually and in combination. We first treated macrophages with the effective concentrations of TLR3i (10 µM) and ox-LDL (10 µg/mL) and found that combined treatment did not dampen phagocytosis any more than the individual treatments (Figure 4D), suggesting that TLR3 and scavenger receptors act along the same pathway.

Next, inspired by a study that demonstrated *Msr1*^*+/-*^ or *Tlr4*^*+/-*^ single heterozygote mice showed no impairment in the phagocytosis of *E. coli*, but double heterozygotes were defective in phagocytosis^41^, we then tested whether using a lower concentration of ox-LDL and TLR3i individually and in combination would result in synergy. When *Tlr4*^*-/-*^ macrophages were treated with 1 µM TLR3i or 1 µg/mL ox-LDL, there was no difference in the phagocytosis of *C. neoformans* compared to untreated cells (Figure 4E). However, when treated with 1 µM TLR3i and 1 µg/mL ox-LDL together, there was a decrease in phagocytosis (Figure 4E), suggesting that these receptors act in synergy along the same pathway.

### *Tlr4*^*-/-*^ macrophages have increased expression of MSR1

Ox-LDL competitively inhibits most scavenger receptors. To try and discern which may be responsible for the phagocytosis of *C. neoformans*, we analysed the surface expression of the scavenger receptors CD36, MAcrophage Receptor with COllagenous structure (MARCO), and MSR1 using flow cytometry. Wildtype and *Tlr4*^*-/-*^ macrophages had a comparable expression of CD36 (Figure 5A). Both cell types expressed very little MARCO (Figure 5B), in line with studies that show that iBMDMs do not express MARCO^42^. Notably, however, MSR1 expression was significantly higher in *Tlr4*^*-/-*^ macrophages compared to wildtype macrophages (Figure 5C), suggesting that the increased phagocytosis of *C. neoformans* observed in *Tlr4*^*-/-*^ macrophages may be due to their increased expression of MSR1.

**Figure 5:**
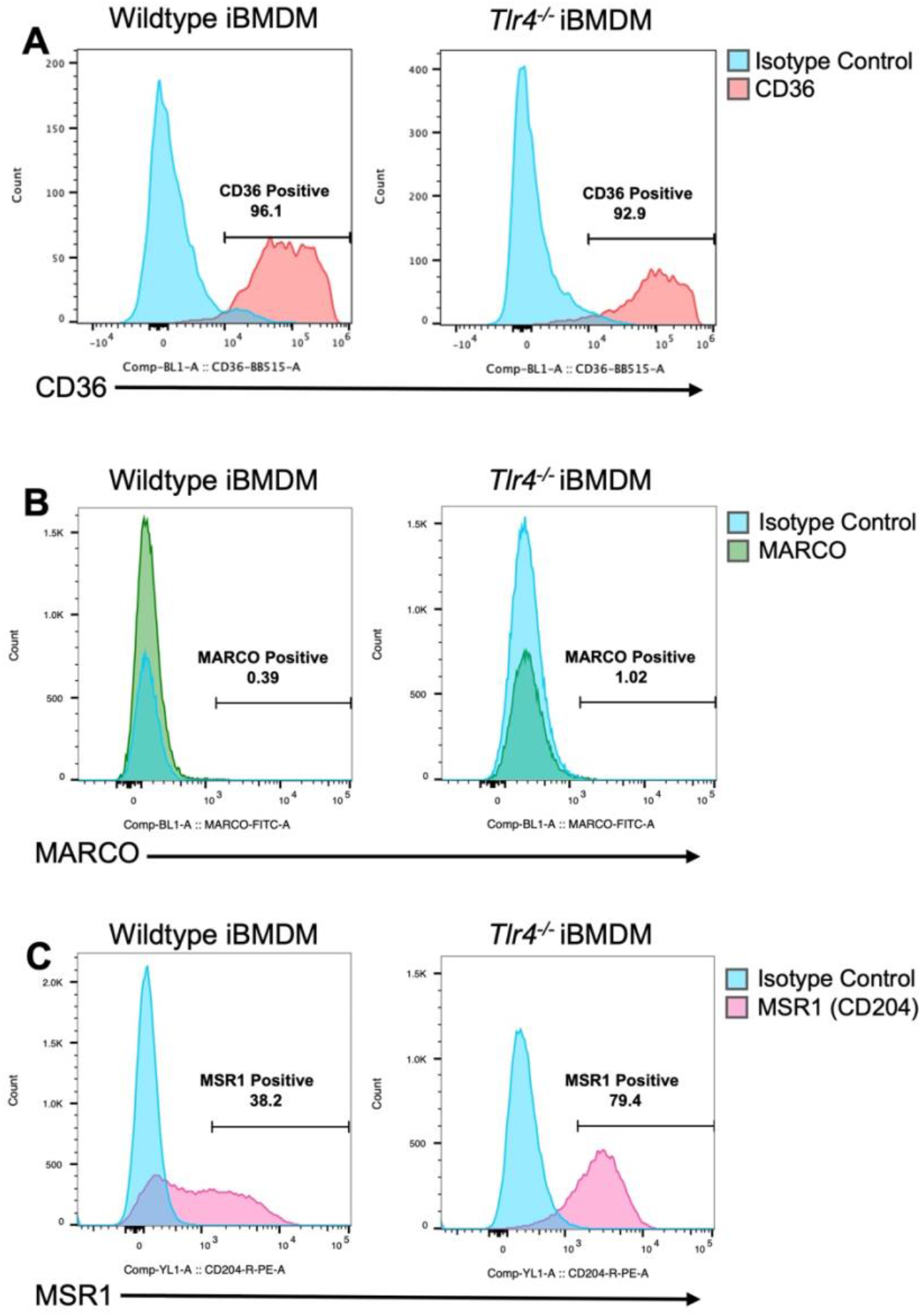
Scavenger receptor expression measured using flow cytometry. Baseline surface expression of **(A)** CD36 stained with anti-mouse CD36-BB515 antibody; **(B)** MAcrophage Receptor with COllagenous structure (MARCO) stained with anti-mouse MARCO-Fluorescein antibody; and **(C)** Macrophage Scavenger Receptor 1 (MSR1), also known as CD204, stained with anti-mouse CD204-PE antibody on wildtype and *Tlr4*^*-/-*^ macrophages. Receptor expression was measured using flow cytometry. Data is representative of three technical repeats.

Given that *MyD88*^*-/-*^ and *Trif*^*-/-*^ macrophages showed a near complete loss of phagocytosis, we hypothesized that these cells express very little MSR1. To test this, the surface expression of MSR1 on these macrophages was also measured using flow cytometry. However, as with *Tlr4*^*-/-*^ macrophages, *MyD88*^*-/-*^ and *Trif*^*-/-*^ cells showed increased MSR1 expression compared to wildtype iBMDMs (Figure 6). Moreover, the proportion MSR1 positive cells in *MyD88*^*-/-*^ and *Trif*^*-/-*^ macrophages was similar to that observed in *Tlr4*^*-/-*^ macrophages (20.4% for wildtype, 71.8% for *Tlr4*^***-/-***^, 65.6% for *MyD88*^***-/-***^ and 74.9% for *Trif*^***-/-***^ macrophages (data not shown)). This implies that increased MSR1 expression alone is not sufficient to drive increased phagocytosis. Either MyD88 and TRIF themselves or some other MyD88- and/or TRIF-dependent molecules may serve as adaptor proteins or coreceptors necessary to drive pathogen engulfment.

**Figure 6:**
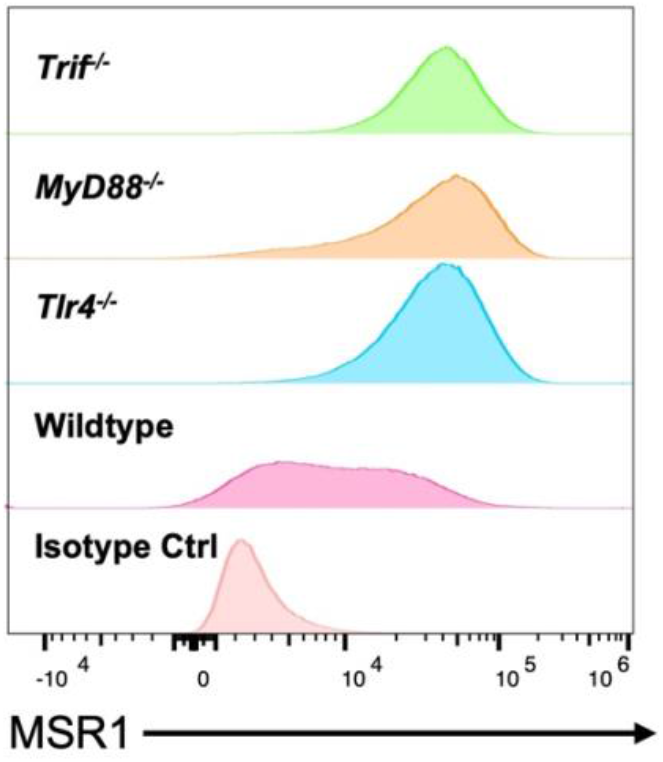
Macrophage Scavenger Receptor 1 (MSR1) expression in wildtype, *Tlr4*^*-/-*^, *MyD88*^*-/-*^ and *Trif*^*-/-*^ macrophages. Baseline surface expression of MSR1 (also known as CD204) was measured using anti-mouse CD204-PE antibody. PE-labelled rat IgG2a kappa was used as an isotype control. Receptor expression was measured using flow cytometry and analysed using the FlowJo software. Data is representative of two technical repeats.

### TLR3 inhibition does not alter the surface expression of MSR1

To further disentangle the interaction between scavenger receptors and TLR3, we measured MSR1 expression in wildtype and *Tlr4*^*-/-*^ iBMDMs after 1 h TLR3 inhibition. We found no significant changes in MSR1 expression after both wildtype and *Tlr4*^*-/-*^ macrophages were treated with the TLR3 inhibitor (Figure 7). We previously showed synergy between ox-LDL pre-treatment and TLR3 inhibition (Figure 4C and 4D); however, this finding suggests that the interaction between MSR1 and TLR3 is not at the level of direct TLR3-mediated regulation of MSR1 expression.

**Figure 7:**
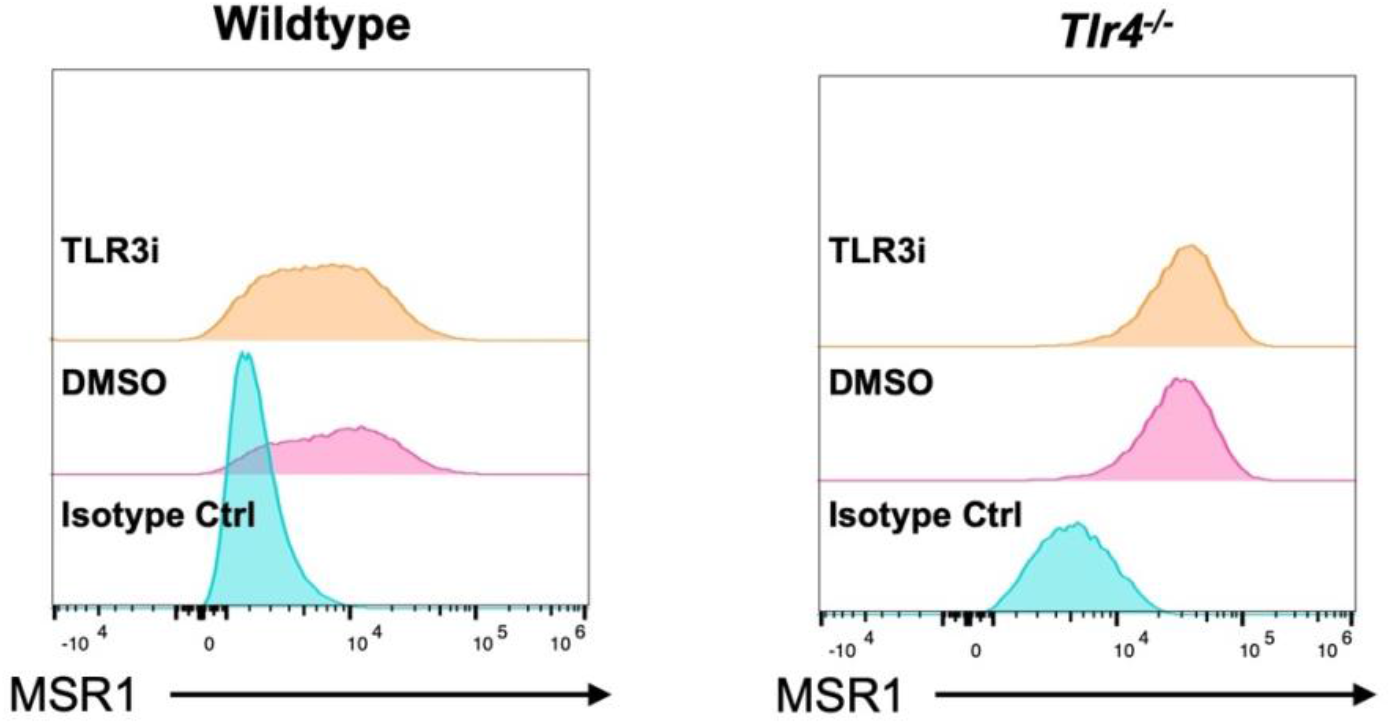
Macrophage Scavenger Receptor 1 (MSR1) expression after TLR3 inhibition. Wildtype and *Tlr4*^*-/-*^ iBMDMs were treated with 0.025% DMSO (control) or 10 µM TLR3/dsRNA complex inhibition. After which MSR1 (also known as CD204) expression was measured by incubating macrophages with anti-mouse CD204-PE monoclonal antibody and running the samples through a flow cytometer. For the isotype control, wildtype and *Tlr4*^*-/-*^ iBMDMs were incubated with rat IgG2a kappa isotype control, PE-labelled.

### MSR1 is a major PRR for the phagocytosis of nonopsonized *C. neoformans*

Having shown that MSR1 is upregulated in *Tlr4*^*-/-*^ macrophages, we next wanted to test the role of individual scavenger receptors in phagocytosis of cryptococci. To achieve this aim, we infected MPI cells (a non-transformed GM-CSF-dependent murine macrophage cell line^42^) derived from wildtype, *Msr1*^*-/-*^, *Marco*^*-/-*^ or double knockout (DKO) C57BL/6 mice with *C. neoformans*. Whilst uptake by *Marco*^*-/-*^ macrophages was indistinguishable from wildtype cells, macrophages derived from *Msr1*^*-/-*^ mice showed a 75% decrease in phagocytosis of nonopsonized cryptococci (Figure 8), suggesting that MSR1 is a critical phagocytic receptor for *C. neoformans*.

**Figure 8:**
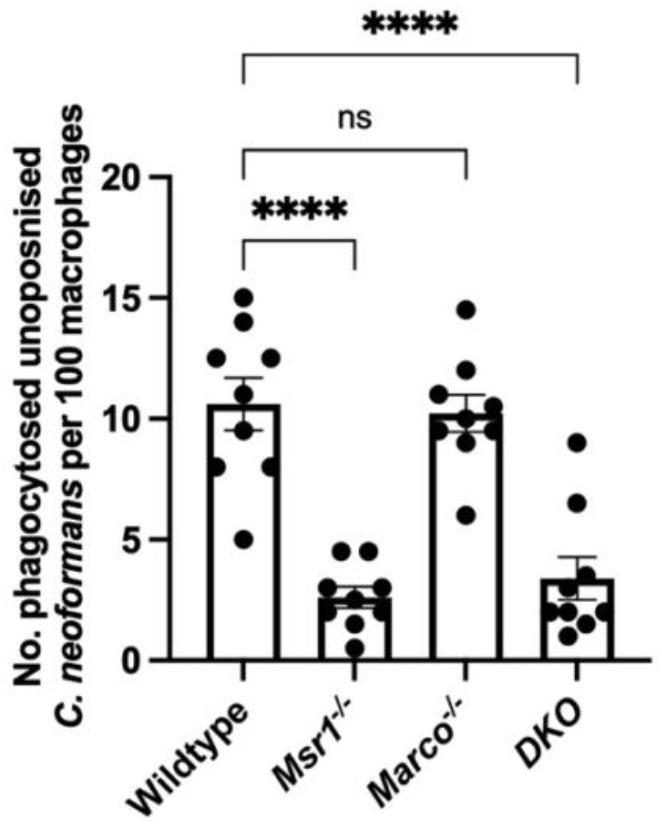
Macrophage Scavenger Receptor 1 (MSR1) is a critical receptor in the phagocytosis of *C. neoformans*. MPI cells, a non-transformed GM-CSF-dependent murine macrophage cell line, were isolated from wildtype, *Msr1*^*-/-*^, *Marco*^*-/-*^ and *MSR1/MARCO* double knockout (DKO) mice, were infected with nonopsonized *C. neoformans*. The data shown is pooled from three technical repeats each performed in triplicates. Data is presented as mean ± SEM; ns, not significant, ****p<0.0001 in a one-way ANOVA followed by Tukey’s post-hoc test.

To provide further support that MSR1 is directly involved in the uptake of *C. neoformans, Tlr4*^*-/-*^ macrophages infected with cryptococci were stained with an anti-MSR1 antibody to investigate the localisation of MSR1 following infection. Immunofluorescence images revealed that MSR1 colocalises with phagocytosed *C. neoformans* (Figure 9, white arrows). Interestingly, not all internalised *C. neoformans* colocalised with MSR1 (Figure 9, green arrows). This suggests specificity of the observed colocalization events and may reflect variation in phagosome maturation stages, such that phagosomes with no MSR1/*C. neoformans* colocalization may have already recycled MSR1 back to the plasma membrane.

**Figure 9:**
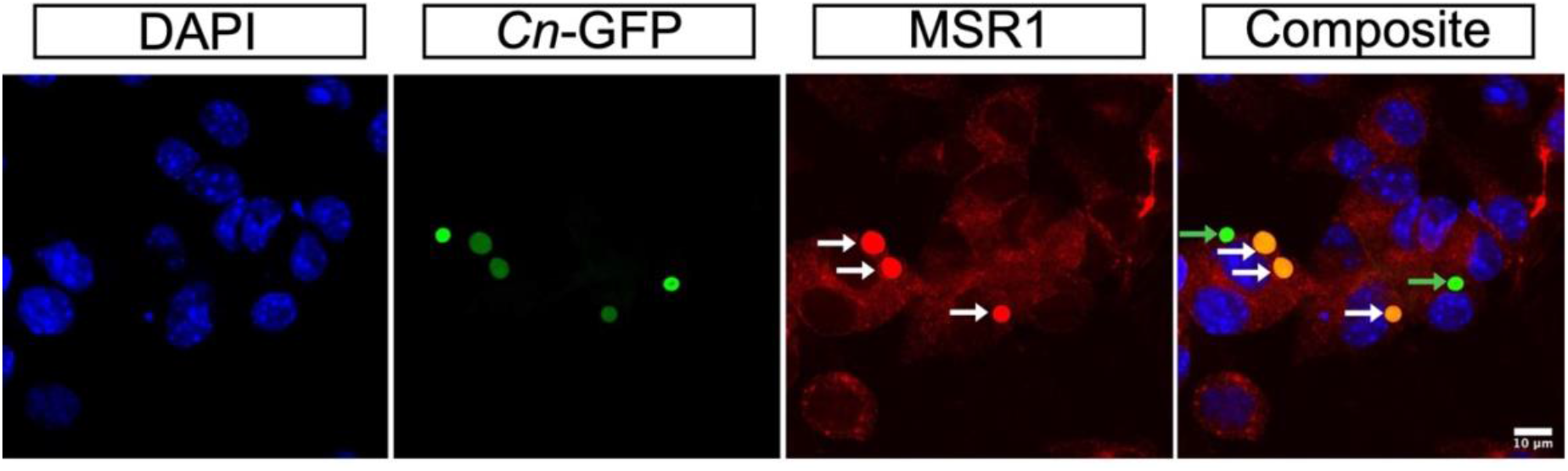
Immunofluorescence analysis of Macrophage Scavenger Receptor 1 (MSR1) expression on *Tlr4*^*-/-*^ macrophages after infection with GFP-expressing *C. neoformans* (*Cn*-GFP). Post infection, macrophages were fixed, permeabilised and stained with rat anti-Mouse CD204 (MSR1) followed by AlexaFluor 594 goat anti-rat IgG secondary antibody. Mounting medium containing DAPI was used to stain the nucleus. Prepared samples were analysed using a confocal microscope at 63X Oil magnification. Representative image shows MSR1 colocalization with phagocytosed *C. neoformans* (white arrows), as well as phagocytosed cryptococci not localised with MSR1 (green arrows). Scale bar = 10 µm

## Discussion

Our findings provide novel insight into the role TLR4 in the phagocytosis of nonopsonized *C. neoformans* by macrophages. We found that the loss of TLR4 signalling unexpectedly increased the phagocytosis of *C. neoformans* by upregulating MSR1 expression. The increase in phagocytosis was driven by crosstalk between TLR4 and TLR3 in a MyD88- and TRIF-dependent manner to modulate the expression of the phagocytic receptor, MSR1. Using *MSR1*^*-/-*^ macrophages we show for the first time that MSR1 is a necessary phagocytic receptor for the uptake of *C. neoformans*. This provides an explanation for the minimal involvement of Dectin-1 in host response to *C. neoformans*^5,43^, emphasizing the significance of our finding.

Scavenger receptors are phagocytic receptors found on the plasma membrane of various immune cells including macrophages^36^. They were first found to bind modified low-density lipoproteins (LDL), but are now known to recognize a wide range of host and microbial ligands such as apoptotic cells, phospholipids, proteoglycan, LPS, and fungal β-glucans^36–38^. It has previously been reported that TLR4 synergises with MSR1 to promote the phagocytosis of Gram-negative *E. coli*, while TLR2 synergises with MSR1 in the phagocytosis of Gram-positive *Staphylococcus aureus*^41^. Similarly, MSR1 was involved in the phagocytosis of the Gram-negative bacteria *Neisseria meningitidis*, which is also recognised by TLR4, while modulating TLR4-mediated inflammatory response to *N. meningitidis* infection^44^. Despite these studies on bacterial pathogens, to our knowledge, ours is the first to report TLR/MSR1 crosstalk in the context of a fungal infection. It is therefore likely that TLR-SR crosstalk to regulate phagocytosis and cytokine expression is a general phenomenon of host-pathogen interactions.

Interestingly, we implicate TLR3, an endosomal PRR known for its role as a dsRNA receptor^45^, in this crosstalk. The mechanism by which TLR4-TLR3 crosstalk regulates the expression and/or activity of MSR1 remains unclear. This is particularly difficult to decipher since most SRs, including MSR1, have very short cytoplasmic tails with no discernible signalling domains^37^. Additionally, since TLR3 is a dsRNA receptor, there is no obvious TLR3 ligand in *C. neoformans*, hence the mechanism driving TLR3 contribution to the modulation of *C. neoformans* uptake by macrophages requires further study. However, our findings suggest a role for MyD88 and TRIF in this TLR4/TLR3/MSR1 axis. Despite the significant decrease in *C. neoformans* phagocytosis observed in *MyD88*^*-/-*^ and *Trif*^*-/-*^ macrophages, we also found that *MyD88*- and *TRIF*-deficient macrophages had increased expression of MSR1. Therefore, their role in the phagocytosis of *C. neoformans* is probably not due to an impact on MSR1 expression. Instead, they may function as coreceptors or activators of some other partner molecule necessary for successful MSR1-mediated pathogen engulfment. The same could be said for TLR3, since treatment with a TLR3 inhibitor had no impact on SRA1 expression even though TLR3 inhibition resulted in decreased phagocytosis.

Others have investigated the role of TLR4 during host response to *Cryptococcus* infection; however, these studies have revealed contradictory results^9,29,46,47^. An *in vitro* study found that the stimulation of microglial cells isolated from the brain of wildtype mice with the TLR4 agonist, lipopolysaccharide (LPS), resulted in increased phagocytosis and killing of *C. neoformans* in a MyD88-dependent manner^48^. Interestingly, *in vivo* studies using *TLR4*-deficient mice have found that the receptor is dispensable during host response to infection^29,30,46^. MyD88 is a key adaptor molecule downstream of all TLRs except TLR3. Mice deficient in *MyD88* consistently show that this adaptor molecule plays a major role in anti-*Cryptococcus* immune response^30,46^, thereby implicating the upstream TLRs in host response. However, to date, the precise role of individual TLRs, including TLR4, during cryptococcal infection is poorly understood. Our data suggest that one possible explanation for these previous conflicting results is varying level of MSR1 expression, which was unaccounted for in these studies. Here we show that the knockout of *TLR4* increased MSR1 expression; however, others have shown that LPS-mediated stimulation of TLR4 was also capable of increasing the expression of scavenger receptors leading to increased uptake^40,49^. Despite expecting TLR4-deficiency and pre-treatment with a TLR4 agonist to have opposing effects, our data implies that TLR can also act as a negative regulator of MSR1, such that any perturbation of TLR4 signalling affects scavenger receptor expression which could impact macrophage response to infection.

It is notable that we find no involvement of MARCO during phagocytosis in our experimental system. It has previously been shown that *Marco*^*-/-*^ mice infected with *C. neoformans* had a significantly higher lung fungal burden compared to control mice^15^. Moreover, alveolar macrophages isolated from *Marco*^*-/-*^ mice had decreased phagocytosis. This is contradictory to our finding that *Marco*^*-/-*^ MPI cells had a comparable level of phagocytosis to wildtype MPI cells. Though this may be explained by the absence of LPS stimulation in our experimental design since others have shown that MARCO expression is inducible by LPS, leading to increased phagocytosis of bacteria^40,50^.

On the other hand, an *in vivo* study using MSR1^-/-^ mice found that knockout mice had reduced lung fungal burden and decreased expression of T-helper 2 (Th2) cytokines, which is an immune polarization state that promotes fungal growth and dissemination^13^. Thus, the authors concluded that MSR1 is normally hijacked by *C. neoformans* to promote its pathogenesis. If this is the case, the increased expression of MSR1 that we observe in *Tlr4*^*-/-*^ macrophages could correlate with poor disease outcome. In support of this idea is the finding that *C. neoformans* clinical isolates that are more readily phagocytosed showed increased brain fungal burden, reduced mice survival and polarization towards the nonprotective Th2 response^51^. Similarly, clinical isolates with low phagocytic indexes were associated with poor fungal clearance (even with antifungal treatment) in the cerebrospinal fluid^52^. Meanwhile, isolates with high phagocytic indexes were associated with increased mortality^52,53^. Therefore, both very high and very low phagocytosis are predictors of poor disease outcome, implying the existence of a ‘Goldilocks’ level of uptake. The more we understand about the clinical outcomes associated with increased phagocytosis compared to decreased phagocytosis of *C. neoformans* will point towards the appropriate way to manipulate MSR1 as a potential therapeutic approach.

In summary, here we present the significance of TLR4/TLR3/MSR1 crosstalk in the phagocytosis of *C. neoformans*, identify MSR1 as a critical receptor for the nonopsonic phagocytosis of *C. neoformans* and support the paradigm that TLRs collaborate with other cell surface receptors to modulate pathogen recognition.

## Materials & Methods

### Tissue Culture and Macrophage Cell Lines

The J774A.1 cell line was cultured in T-75 flasks [Fisher Scientific] in Dulbecco’s Modified Eagle medium, low glucose (DMEM) [Sigma-Aldrich], containing 10% live fetal bovine serum (FBS) [Sigma-Aldrich], 2 mM L-glutamine [Sigma-Aldrich], and 1% Penicillin and Streptomycin solution [Sigma-Aldrich] at 37°C and 5% CO_2_. During phagocytosis assays, J774A.1 macrophages were seeded at a density of 1×10^5^ cells per well of a 24-well plate [Greiner Bio-One].

Immortalised bone marrow derived macrophages were originally isolated from C57BL/6 wildtype, *Tlr4*^*-/-*^, *MyD88*^*-/-*^ and *Trif*^*-/-*^ single knock out mice and immortalised via transformation with retroviruses expressing Raf and Myc, two well-known proto-oncogenes^54^. Immortalised BMDMs were cultured in DMEM, low glucose [Sigma-Aldrich] supplemented with 10% heat inactivated FBS [Sigma-Aldrich], 2 mM L-glutamine [Sigma-Aldrich], and 1% Penicillin and Streptomycin solution [Sigma-Aldrich] at 37°C and 5% CO_2_. During phagocytosis assays, iBMDMs were seeded at a density of 3×10^5^ cells per well of a 24-well plate [Greiner Bio-One].

Max Plank Institute (MPI) cells are a non-transformed, granulocyte-macrophage colony-stimulating factor (GM-CSF)-dependent murine macrophage cell line that is functionally similar to alveolar macrophages^42,55^. In this study, MPI cells from wildtype, *Msr1*^*-/-*^, macrophage receptor with collagenous structure knockout (*Marco*^*-/-*^) and *MSR1/MARCO* double knockout (DKO) C57BL/6 mice were utilised. Cells were cultured in Roswell Park Memorial Institute (RPMI) 1640 medium [ThermoFisher] supplemented with 10% heat inactivated FBS [Sigma-Aldrich], 2 mM L-glutamine [Sigma-Aldrich], and 1% Penicillin and Streptomycin solution [Sigma-Aldrich] at 37°C and 5% CO_2_. Each flask was further supplemented with 1% vol/vol GM-CSF conditioned RPMI media prepared using the X-63-GMCSF cell line. When being used in phagocytosis assays, MPI cells were seeded at a density of 2×10^5^ cells per well of a 24-well plate [Greiner Bio-One] with 1% vol/vol GM-CSF.

### Phagocytosis Assay

Phagocytosis assays were performed to measure the uptake of *Cryptococcus* by macrophages under various conditions. Twenty-four hours before the start of the phagocytosis assay, the desired number of macrophages were seeded onto 24-well plates in complete culture media. The cells were then incubated overnight at 37°C and 5% CO_2_. At the same time, an overnight culture of *Cryptococcus neoformans* var. *grubii* KN99α strain, that had previously been biolistically transformed to express green fluorescent protein (GFP)^56^, was set up by picking a fungal colony from YPD agar plates (50 g/L YPD broth powder [Sigma-Aldrich], 2% Agar [MP Biomedical]) and resuspending in 3 mL liquid YPD broth (50 g/L YPD broth powder [Sigma-Aldrich]). The culture was then incubated at 25°C overnight under constant rotation (20rpm).

On the day of the assay, macrophages were activated using 150ng/mL phorbol 12-myristate 13-acetate (PMA) [Sigma-Aldrich] for 1 h at 37°C. PMA stimulation was performed in media containing heat-inactivated serum (iBMDMs and MPI cells) or in serum-free media (J774A.1) to eliminate the contribution of complement proteins during phagocytosis. Where applicable, macrophages were then treated with the desired concentration of soluble inhibitors of PRRs (Table 1) and incubated at 37°C for 1 h. Meanwhile pre-treatment with the general scavenger receptor ligand, oxidised low-density lipoprotein (ox-LDL), occurred for 30 mins. The concentration used for each molecule is indicated in Table 1 and in the corresponding results.

**Table 1:** List of inhibitors and ligands used to block macrophage pattern recognition receptor (PRRs) activity.

| Molecule Name | Target Receptor | Concentration(s) Used | Mode of Action | Cat# [Brand] |
| --- | --- | --- | --- | --- |
| TAK-242 | TLR4 | 0.2 μM | Selectively binds to Cys747 of TLR4 and disrupts its interaction with the adaptor molecules, TIRAP and TRAM. | 614316 [Sigma-Aldrich] |
| CU CPT 22 | TLR2/1 | 1 μM | Competes with lipoproteins for binding to TLR2/1 and inhibits the release of proinflammatory cytokines TNF-α and IL-1β. | 4884 [Tocris] |
| ODN-2088 | TLR9 | 1 μM | An inhibitory sequence that disrupts the colocalization of CpG oligonucleotides with TLR9 in the endosome | tlrl-2088 [InvivoGen] |
| TLR3/dsRNA Complex Inhibitor | TLR3 | 1 μM – 10 μM | A specific and competitive inhibitor of dsRNA binding to TLR3 | 614310 [Sigma-Aldrich] |
| Pepinh-MYD | MyD88 | 20 μM | A peptide that inhibits the homodimerization of MyD88 | tlrl-pimyd [InvivoGen] |
| Pepinh-TRIF | TRIF | 20 μM | An inhibitory peptide that obstructs TLR-TIF interaction. | tlrl-pitrif [InvivoGen] |
| B1605906 | IKKβ | 5 μM | Binds to IKKβ and prevents the activation of NF-κB. | A kind gift from Dario Alessi, University of Dundee |
| MRT67307 | TBK1 | 10 μM | Blocks the activity of TBK1 and IKKε thereby preventing the activation of IRF3. | A kind gift from Dario Alessi, University of Dundee |
| Low Density Lipoprotein from Human Plasma, oxidised (ox-LDL) | Scavenger Receptors | 10 μg/mL | A known scavenger receptor ligand | L34357 [ThermoFisher] |

To prepare *C. neoformans* for infection, the overnight *C. neoformans* culture was washed two times in 1X PBS and centrifuged at 6500 rpm for 2.5 mins. To infect macrophages with nonopsonized *C. neoformans*, after the final wash, the *C. neoformans* pellet was resuspended in 1 mL PBS, counted using a hemacytometer, and fungi incubated with macrophages at a multiplicity of infection (MOI) of 10:1. The infection was allowed to take place for 2 h at 37°C and 5% CO_2_. Infection occurred in the presence of soluble inhibitors.

In some instances, macrophages were infected with antibody-opsonized *C. neoformans*. To opsonize the fungi, 1×10^6^ yeast cells in 100 μL PBS were opsonized for 1 h using 10 μg/mL anti-capsular 18B7 antibody (a kind gift from Arturo Casadevall, Albert Einstein College of Medicine, New York, NY, USA). After 2 h infection, macrophages were washed 4 times with PBS to remove as much extracellular *C. neoformans* as possible.

### Fluorescent Microscopy Imaging

Having washed off extracellular cryptococci, the number of phagocytosed fungi was quantified using images from a fluorescent microscope. To distinguish between phagocytosed and extracellular *C. neoformans*, wells were treated with 10 μg/mL calcofluor white (CFW) [Sigma-Aldrich], a fluorochrome that recognises cellulose and chitin in cell walls of fungi, parasite and plants^57^, for 10 mins at 37°C. Next, fluorescent microscopy images were acquired using the Zeiss Axio Observer [Zeiss Microscopy] fitted with the ORCA-Flash4.0 C11440 camera [Hamamatsu] at 20X magnification. The phase contrast objective, EGFP channel and CFW channel were used. Image acquisition was performed using the ZEN 3.1 Blue software [Zeiss Microscopy] and the resulting images were analysed using the Fiji image processing software [ImageJ].

To quantify the number of phagocytosed cryptococci, the total number of ingested *C. neoformans* was counted in 200 macrophages, then the values were applied to the following equation: ((number of phagocytosed *C. neoformans*/number of macrophages) * 100). Therefore, the result of the phagocytosis assay is presented as the number of internalised fungi per 100 macrophages.

### Live Imaging

To assess the intracellular proliferation rate (IPR) of *C. neoformans* within macrophages, infected macrophages were captured at a regular interval over an extended period. Live-cell imaging was performed by running the phagocytosis assay as usual, then after washing off extracellular cryptococcus, the corresponding media for the macrophage cell line was added back into the well before imaging. Live imaging occurred using the Zeiss Axio Observer at 20X magnification and images were acquired every 5 mins for 18 hours at 37°C and 5% CO_2_.

The resulting videos were analysed using Fiji [ImageJ] and IPR was determined by quantifying the total number of internalised fungi in 200 macrophages at the ‘first frame’ (time point 0 (T0)) and ‘last frame’ (T10). Then, the number of phagocytosed fungi at T10 was divided by the number of phagocytosed fungi at T0 to give the IPR (IPR = T10/T0).

### Immunofluorescent Imaging

Immunofluorescence was used to investigate receptor localisation on macrophages. Firstly, 13 mm cover slips were placed onto 24-well plates prior to seeding with desired number of macrophages. After overnight incubation, macrophages were used in a standard phagocytosis assay. Prior to staining, macrophages were fixed with 4% paraformaldehyde for 10 mins at room temperature and permeabilised with 0.1% Triton X-100 diluted in PBS for 10 mins at room temperature. To stain for MSR1 localisation, 5 µg/mL rat anti-mouse CD204 (MSR1)-PE [Fisher Scientific; Cat#: 12-204-682] was used as the primary antibody. Cells were incubated with the primary antibody for 1 h at room temperature. After washing twice with PBS, macrophages were incubated with 5 µg/mL AlexaFluor 594 goat anti-rat IgG secondary antibody [ThermoFisher Scientific; Cat#: A11007] for 1 h at room temperature in the dark. Coverslips were mounted on 5 µL VECTASHIELD HardSet antifade mounting medium with DAPI [Vector Laboratories]. Images were acquired using the Zeiss LSM900 Confocal with Airyscan2, laser lines 405, 488, 561 and 640 nm, and at 63X oil magnification. Image acquisition was performed using the ZEN 3.1 Blue software [Zeiss Microscopy] and the resulting images were analysed using the Fiji image processing software [ImageJ].

### Flow Cytometry

Flow cytometry was used to measure the surface expression of scavenger receptors on macrophages. Prior to staining, macrophages were incubated with 2.5 µg/mL rat anti-mouse CD16/CD32 Fc block [BD Biosciences; Cat#: 553142] diluted in FACS buffer (1XPBS without Mg^2+^ and Ca^2+^ supplemented with 2% heat inactivated FBS and 2 mM EDTA). After Fc blocking, the desired concentration of fluorochrome-conjugated antibodies diluted in FACS buffer was added into each tube still in the presence of the Fc block mixture. The following fluorochrome-conjugated antibodies were used: 0.5 µg/mL anti-mouse CD45-PerCP-Cyanine5.5 [ThermoFisher; Cat#: 45-0451-82], 0.25 µg/100 µL anti-mouse CD204(MSR1)-PE [Fisher Scientific; Cat#: 12-204-682], 0.25 µg/100 µL anti-mouse CD36-BB515 [BD Biosciences; Cat#: 565933], and 10 µL/100 µL anti-mouse MARCO-Fluorescein [Biotechne; Cat#: FAB2956F]. Fluorescent minus one (FMO) controls were included to aid in setting gating boundaries. Isotype controls were used to test for non-specific binding. The following isotype control antibodies were used: 0.25 µg/100 µL PE rat IgG2a, κ isotype control [Fisher Scientific; Cat#: 15248769], 0.25 µg/100 µL BB5151 Mouse IgA, κ isotype control [BD Biosciences; Cat#: 565095], and 10 µL/100 µL Rat IgG_1_ Fluorescein isotype control [Biotechne; Cat#: IC005F]. Finally, samples stained with only one fluorophore were used as compensation controls. After staining, samples were resuspended in FACS buffer for single staining and unstained controls and FACS buffer with DAPI [ThermoFisher], a live dead stain, for all other samples.

Stained samples were run on the Attune NxT flow cytometer [ThermoFisher] and acquired using the Attune NxT software [ThermoFisher]. The resulting data was analysed using the FlowJo v10 software [BD Life Sciences]. Before determining the proportion of macrophages positive for a particular fluorochrome, a gating strategy was employed to achieve the sequential exclusion of debris and doublets (Supplementary Figure 2). Anti-CD45-PerCP-Cyanine5.5 was used to identify total leukocytes, and DAPI was used to exclude dead cells.

### Statistics

GraphPad Prism Version 9 for Mac (GraphPad Software, San Diego, CA) was used to generate graphical representations of experimental data. Inferential statistical tests were performed using Prism. The data sets were assumed to be normally distributed based on the results of a Shapiro-Wilk test for normality. Consequently, to compare the means between treatments, the following parametric tests were performed: unpaired t test, one-way ANOVA, and two-way ANOVA. ANOVA tests were followed up with Tukey’s post-hoc test. Variation between treatments was considered statistically significant if p-value < 0.05.

## Acknowledgements

We thank Dr. Maria Makarova for assistance with confocal microscopy. The *MSR1* knockout and *MSR1/MARCO* double knockout lines were established with the support of NC3Rs Grant NC/V001019/1. Work in C.E.B lab is supported by grant BB/V000276/1. This work was partly supported by a Royal Society project grant (RGS\R2\202260) to S.M lab. C.U.O is supported by a PhD studentship from the Darwin Trust of Edinburgh. A.L.W. was supported by PhD funding from the Wellcome Trust ‘MIDAS’ doctoral training program. R.C.M. gratefully acknowledges support from the BBSRC and European Research Council Consolidator Award.

## Author Contributions

R.C.M conceived the project. A.L.W and G.D. generated preliminary data that influenced the conception of the project. C.U.O and R.C.M generated hypotheses and designed the experiments. C.U.O performed all the experiments, analysed the data and wrote the manuscript. G.D. helped perform the flow cytometry experiments. C.E.B provided *Tlr4*^*-/-*^, *MyD88*^*-/-*^, *Trif*^*-/-*^ immortalised BMDMs. S.M, S.G, and G.F provided *Msr1*^*-/-*^, *Marco*^*-/-*^ and *DKO* MPI cells. O.D.C provided transfected cell lines that contributed to hypothesis generation. S.M and S.G provided critical discussion of the experimental results. All authors contributed to the review of the manuscript. R.C.M acquired funding.

## Competing interests

The authors declare no competing interests.

**Supplementary Figure 1:**
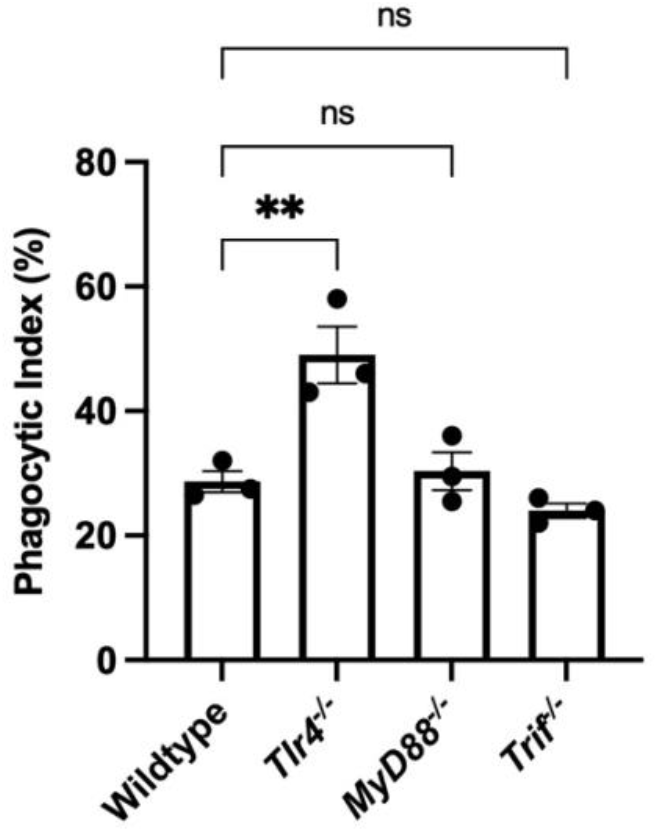
*Candida albicans* infection of immortalised bone marrow derived macrophages (iBMDMs). iBMDMs isolated from wildtype, *Tlr4*^*-/-*^, *MyD88*^*-/-*^ and *Trif*^*-/-*^ mice were infected with CAF2-1 *Candida albicans* expressing dTomato fluorescent protein (Caf2-dTomato) at a multiplicity of infection of 5 *C. albicans* to 1 macrophage for 45 mins. Phagocytosis was quantified as the percentage of macrophages that internalised at least one fungus, termed phagocytic index. Data is representative of two technical repeats each performed with triplicate biological repeats. Data shown as mean ± SEM; ns, not significant, **p<0.01 in a one-way analysis of variance (ANOVA) followed by Tukey’s post-hoc test.

**Supplementary Figure 2:**
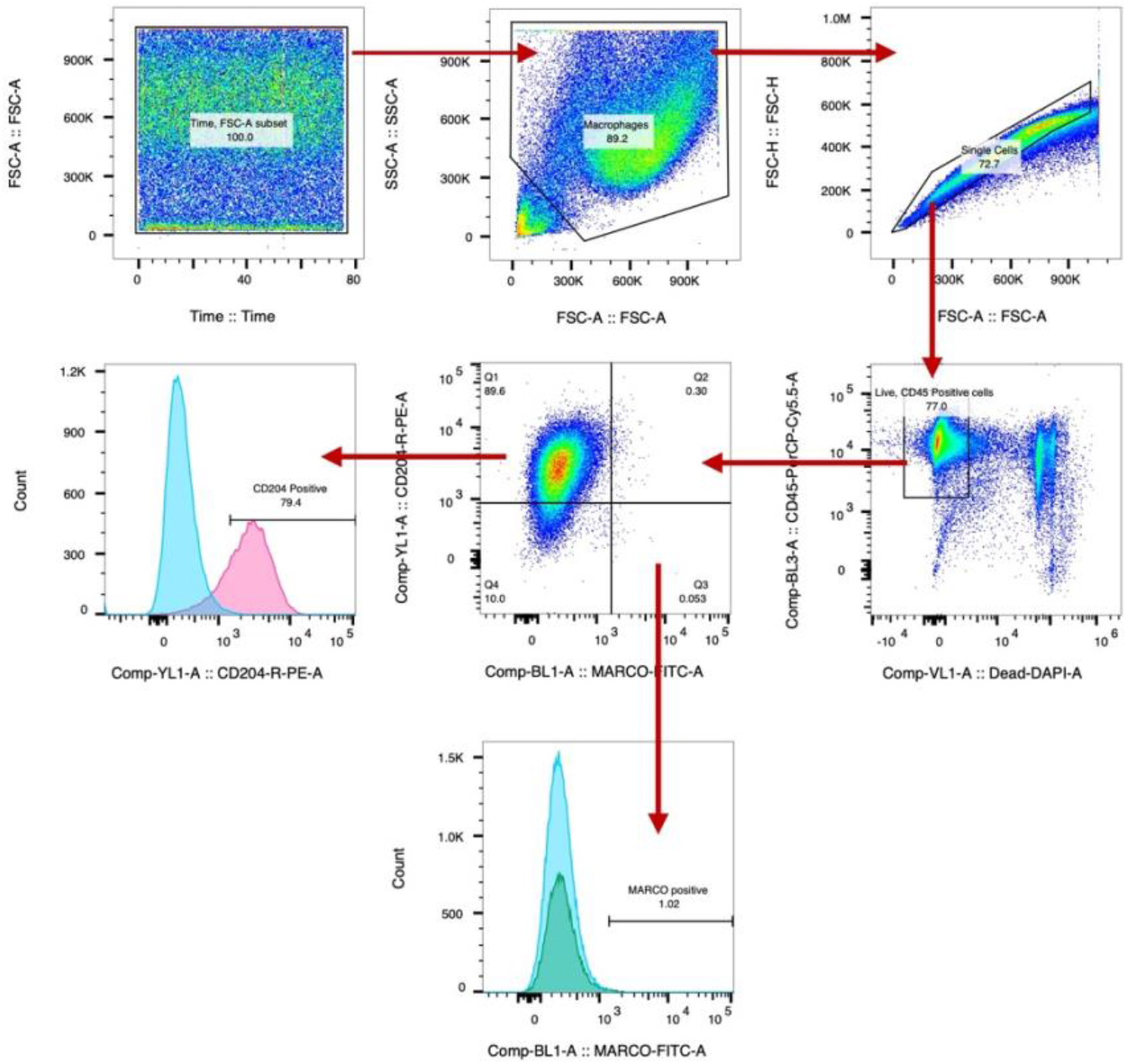
Flow cytometry gating strategy to detect scavenger receptor positive populations. The above example shows *Tlr4*^*-/-*^ macrophages stained with anti-CD204, PE conjugated and anti-MARCO, FITC conjugated antibodies. Following macrophage staining with relevant antibodies, cells were run through the Attune NxT flow cytometer and data acquired using the Attune NxT software. Data analysis was performed using FlowJo v10 software. Firstly, a time gate was applied to examine the quality of the sample run. Next, gates were applied to exclude debris and doublets. Anti-CD45-PerCP-Cyanine5.5 was used to identify total leukocytes, and DAPI was used to exclude dead cells. From the resulting population of live CD45+ cells, gates were applied to detect CD204 (MSR1) and MARCO scavenger receptor positive populations.

